# Motor control processes moderate visual working memory gating

**DOI:** 10.1101/2025.03.05.641641

**Authors:** Şahcan Özdemir, Eren Günseli, Daniel Schneider

**Author notes:** Corresponding Author: Şahcan Özdemir.

## Abstract

Gating processes that regulate sensory input into visual working memory (WM) and the execution of planned actions share neural mechanisms, suggesting a mutual interaction. In a preregistered study (OSF), we examined how this interaction may result in sensory interference during WM storage using a delayed-match-to-sample task. Participants (12 male, 20 female) memorized the color of a target stimulus for later report on a color wheel. The shape of the target indicated which hand they would adjust the color wheel with. During the retention interval, an interference task was presented, requiring a response with either the same or different hand as the main task. In half of the interference trials, the interfering task cue was also colored to introduce visual interference. EEG results showed early motor planning during sensory encoding, evidenced by mu/beta suppression contralateral to the responding hand. The interference task only impaired WM performance when it included an irrelevant color, indicating that the interference effect was primarily driven by the irrelevant sensory information. In addition, color reporting in the WM task was biased toward the irrelevant color. This was more pronounced when both tasks were performed with the same hand, suggesting a selective gating mechanism dependent on motor control processes. This effect was mitigated by a control mechanism, which was evident in frontal theta activity, where higher power predicted lower bias on the single-trial level. Our findings thus reveal that sensory WM updating can be induced by interfering motor actions, which can be compensated by a reactive control mechanism.

**SIGNIFICANCE STATEMENT:** Working memory is increasingly recognized not just as a passive information storage but as an active mechanism that constructs prospective representations to guide future actions. We investigated how future-oriented plans regulate the entry of new information for maintenance. We found that when a stored memory is linked to a response, it becomes particularly vulnerable to interference from sensory input that demands the same response. We also identified neural signatures of this interaction where a control mechanism mitigates interference from irrelevant information. These findings provide key insights into the fundamental architecture of memory, demonstrating for the first time that prospective motor codes not only shape the use of stored information but also influence how new information is integrated into working memory.

## INTRODUCTION

Working memory (WM) serves a critical function in human cognition by maintaining and manipulating task-relevant information (Baddeley, 2003). Recent research has increasingly focused on the ‘task-relevant’ aspect of WM, emphasizing its prospective nature (Gunseli et al., 2014; Huang, 2015; for reviews see Miller et al., 2018; Olivers & Roelfsema, 2020; Postle, 2006; van Ede, 2020; Van Ede & Nobre, 2023). The shift towards a prospective view has expanded the examination of how sensory representations are maintained and manipulated in a bidirectional relationship with prospective action representations (Boettcher et al., 2021; Nasrawi et al., 2023; Olivers & Roelfsema, 2020; Rösner et al., 2022; Sahakian et al., 2025; Schneider et al., 2017; Trentin et al., 2023). Building on this framework, the present study investigates the influence of action planning on WM gating processes, specifically examining the extent to which the storage of relevant information in WM is susceptible to interfering stimuli.

The prospective view on WM highlights the relationship between sensory and motor representations by showing concurrent preparation of motor plans for prioritized WM items (Boettcher et al., 2021; Nasrawi et al., 2023; Rösner et al., 2022; Schneider et al., 2017; van Ede et al., 2019). In this context, evidence for motor preparation is characterized by the suppression of mu/beta oscillatory power (8-14/14-30 Hz, respectively) over the sensorimotor cortex contralateral to the responding hand, which can occur concurrently with the attentional selection of the target item. And this concurrent motor process affects how information is stored in WM. For example, action selection can influence which memory feature is prioritized (Heuer et al., 2020) or mediate the level of inter-item biases when multiple items in WM are associated with different actions (Trentin et al., 2024).

If motor processes influence how information is stored in WM, might they also regulate how new information enters WM during ongoing storage? Such a relationship is plausible, given evidence of overlapping neural mechanisms between motor control processes and WM input gating, i.e., the regulation of information flow into WM (Chatham & Badre, 2015). Therefore, the planning or execution of a motor response might indirectly influence WM’s susceptibility to new information, either enhancing (open gate) or reducing (closed gate) information uptake. Our investigation specifically examines how this input gating process is modulated by correspondence between motor representations maintained in WM and motor responses executed during an interfering task. We hypothesize that sensory inputs associated with corresponding motor responses are more likely to gain access to WM, suggesting a selective gating mechanism that prioritizes information aligned with stored motor plans.

To test this hypothesis, we employed a delayed match-to-sample WM task that required reporting the color of a target stimulus (Figure 1). We manipulated motor planning by assigning the shape of the target object to indicate the use of either the left or right hand for color report. Additionally, we introduced an interference task during the WM storage interval, requiring participants to perform a secondary action with either the same or a different hand as used in the primary task. In half of the interference trials, the cue during this secondary task was colored, allowing us to introduce sensory interference.

**Figure 1.**
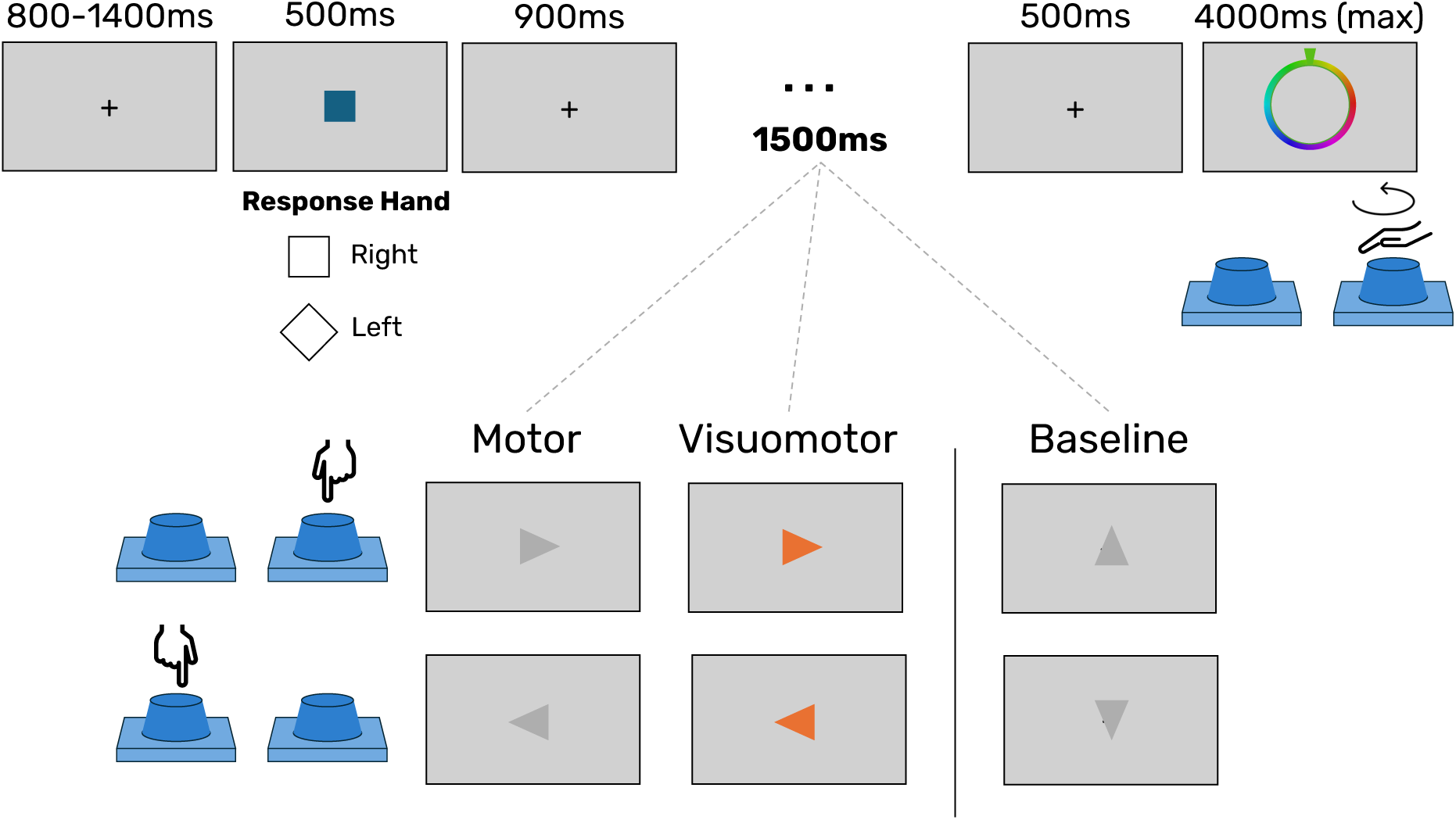
Illustration of the experimental procedure. Participants completed a delayed match-to-sample task, and the response hand was manipulated. The color of the target had to be remembered for later report, while its shape indicated which hand to use for that. Color report was done by adjusting a color wheel to display the target color through rotating one of two response knobs. During the maintenance phase, participants performed an interference task requiring a response to the direction of a central cue (pressing the left vs. right knob) that either corresponded or did not correspond to the hand used in the main task. In half of the interference trials, the interference cue was colored, introducing visual interference. In the baseline condition (cue pointing up or down), there was no response required during maintenance.

Our study was guided by three preregistered hypotheses (OSF). First, we observed the onset of oscillatory correlates of motor planning early during encoding (Hypothesis I), however there was no relation to later retrieval performance. Moreover, we showed higher error rates after visuomotor interference, with no difference between baseline (no interference) and motor interference conditions, indicating that the irrelevant color interfered with WM (Hypothesis II). Finally, we observed a higher attraction bias in WM towards the interference color when the same hand was used for both tasks (Hypothesis III). This effect was accompanied by an increase in frontal theta power following the response to the secondary task, potentially reflecting a reactive control mechanism to reduce the impact of task-irrelevant information on WM (Rac-Lubashevsky & Kessler, 2015, 2018).

## METHODS

### Participants

A total of 32 (M_age_= 23.71; SD_age_ = 2.93; 20 Female) participants participated in the experiment, and were compensated with either money (12€ per hour) or course credits. Two participants were excluded from further analysis due to their misunderstanding of the instructions (i.e., pressing the response buttons already during target presentation), and one participant was excluded based on the exclusion criteria detailed below. Additionally, one participant’s data was used only for the behavioral analysis due to excessive noise in the EEG signal. Thus, the final behavioral analysis included 29 participants while the EEG analyses were conducted with 28 participants.

We calculated the target number of participants using G*Power software (Faul et al., 2007). A priori sample size calculations were conducted for a one-tailed paired-samples t-test referring to the response correspondence effect on attraction bias (Hypothesis III; see Behavioral Analysis section). These calculations employed an alpha level of .05 to control for Type I error, with a desired power of 95%. Given the novelty of the effect under investigation and the lack of directly comparable studies, we used the effect size (Cohen’s d = 0.71) from a study on active vs. passive engagement with a distractor during working memory storage as a reference (Saito et al., 2023). This analysis yielded 23 as the minimum number of participants to achieve the desired statistical power. Since this study did not align perfectly with our research questions, the target number of participants has been increased slightly to account for a potentially weaker effect.

To further ensure the robustness of our sample size, we reviewed studies measuring mu/beta suppression during motor planning in a WM experiment, where three studies used a sample size of 24-26 (Boettcher et al., 2021; Rösner et al., 2022; van Ede et al., 2019). To accommodate the counterbalancing of the assignment of stimulus shape to responding hand (see Procedure), we determined an even number of 26 participants as the target sample (see OSF preregistration). Our final data collection slightly exceeded this target by two datasets (i.e., 28 datasets including EEG data) due to inaccurate coordination with the laboratory staff. Importantly, conducting the analyses only with the first 26 usable datasets did not alter the result patterns.

Data quality is assured by the preregistered exclusion criteria. Participants with an accuracy lower than 70% in using the correct hand in the main task were excluded from further analyses. Furthermore, participants whose mean absolute error was more than 2 standard deviations away from the sample mean were discarded. As indicated above, this led to the rejection of one dataset from all further analyses.

### Ethics statement

This study was conducted under the ethics approval of the local ethics committee at the Leibniz Research Centre for Working Environment and Human Factors and in accordance with the Declaration of Helsinki.

### Apparatus, stimuli, and procedure

The experiment was conducted using a ViSaGe MKII Stimulus Generator (Cambridge Research Systems, Rochester, UK) and presented on a 22-inch CRT screen (resolution: 1024x768, refresh rate: 100 Hz) in a dimly lit, electrically shielded experiment room. The paradigm was prepared and coded on Lazarus IDE with free Pascal. Responses were collected via two response knobs. These knobs were constructed from 3D-printed material, covering two rotary encoders attached to an Arduino processor (Arduino, Lombardia, Italy). The knobs could be rotated continuously, and the rotational data were decoded by the two rotary encoders and transmitted to the experimental setup using Arduino. The minimum step size of the encoder rotation corresponds to 2° on the color wheel. The experiment consisted of 1200 trials in 10 blocks. There was a short break of ∼2 minutes between the blocks and a longer break of ∼5-10 minutes halfway through the experiment.

For stimulus presentation, 180 colors were sampled from HSV color space. The saturation and the value were kept constant (S=.85, V=.9). According to the gradual change in hue (from 1 to 360, 2 degrees per step), 180 colors were sampled at equal distances. Each sample color was used as the target color 6 to 7 times. The sampled 180 colors were then placed on a visual circle to prepare a color wheel (the probe) in each trial. The orientation of the color wheel was randomized to prevent a re-coding of color into a spatial location on the color wheel.

Before the experiment, all participants completed a handedness questionnaire (Oldfield, 1971) to assess their dominant hand. All participants were classified as right-handed. Due to the nature of the experimental task, their color vision was evaluated via the Ishihara Test for Color Blindness.

Participants were asked to complete a delayed match-to-sample WM task in each trial (see Figure 1). First, they were shown a target item centered on the screen to remember its color until the end of the trial. The target was presented as either a square or a diamond shape (1.83° x 1.83°) at the center of the screen. The shape of the target item informed participants whether to use their right or left hand for color report (e.g., right hand for square, left hand for diamond). Shape-hand assignment was counterbalanced across participants. The target presentation was followed by a fixation cross (0.44° x 0.44°) for 900ms. Then, participants were presented with the interference period for 1500ms. What participants encountered in the interference period differed across three conditions: motor interference, visuomotor interference, and baseline.

In the motor interference condition (400 trials), we presented a gray arrow as a cue which either pointed to right or left (HSV: 0, 0, 0.5; side length: 1.83°). Participants were instructed to click the response knob on the side the arrow pointed to as quickly as possible. In the visuomotor interference condition (400 trials) participants were instructed to do the same task, but this time the cue color varied. The cue color was 60-90 degrees away from the target color on the color wheel. The level of difference was preferred to maximize the interference effect (Saito et al., 2023). Thus, the color of the cue added a visual interference aspect to this condition. The varying colors of the visuomotor interference did not create an Oddball-like effect (Polich & Margala, 1997; Reed et al., 2022), as shown by the comparison of the event-related potential (ERP) between the visuomotor and motor interference conditions (Text S2; Figure S2). In the baseline condition (400 trials), participants were shown a gray arrow which pointed either upwards or downwards. In this condition, they were expected to ignore the arrow and respond only to the presentation of the later memory probe. All conditions were presented in a randomly interleaved manner throughout all trials. Regardless of the condition or participants’ response times to the interference task, the cues remained on the screen for 1500ms.

Following this interference period, participants were presented with another fixation cross for 500ms. Then they were instructed to indicate the memorized color on a color wheel (diameter: 5°; thickness: 1°) by rotating the response knob which had been indicated by the shape of the target item at the beginning of the trial. By rotating the knob, participants moved a small arrow (a trapezoid, height: 0.95°; bottom 0.38°; top: 0.23°) over the color wheel to select the desired color. The color of the arrow (or cursor) dynamically updated as it moved along the wheel, matching the currently selected color to facilitate accurate selection. To finalize their response, participants were required to press the knob after having completed their color adjustment. Each trial allowed participants a maximum of three seconds to complete their response. If the participant did not press the knob within this time frame, the last color indicated by the cursor was automatically recorded as the given response.

### Behavioral Analyses

#### Registered analysis

Only trials in which participants responded to both the main task and the interference task using the cued hand were included in the behavioral analysis. In each trial, the absolute difference between the given response and the target color was calculated as an error (relative to their position on the color wheel).

For all behavioral analyses, we provide both frequentist and Bayesian statistics. In Bayesian repeated measures ANOVAs, the BF₁₀ value represents the comparison of the tested model (with a given main effect) against the null model. When analyzing interactions, the null model includes the main effects (Bergh et al., 2020).

For Hypothesis II (see above), absolute errors were averaged for each condition to reveal the magnitude of the interference effect. We expected to see more interference when the interfering task had an irrelevant visual feature (visuomotor condition), compared to the motor interference condition. To test the difference between different types of interference and their relation to the baseline condition, we conducted a one-way ANOVA with three levels (visuomotor interference, motor interference, baseline). All p-values from the multiple comparisons following the ANOVA were Holm adjusted.

Hypothesis III focused on how the WM representation is biased by the interfering visual object in the visuomotor interference condition. To examine this effect, we calculated the attraction bias. Since the effect is calculated relative to the interference color, this could only be realized in the visuomotor interference condition. In each trial, an attraction bias was assumed if the reported color lay in the direction of the interference color on the color wheel. The opposite case is considered a repulsion bias (Chunharas et al., 2022). For analyzing this bias, each error value between the actual and the reported color was assigned a positive or a negative value, depending on whether there was a shift in the direction of or against the position of the interference color. Then, we calculated the average of assigned errors for each condition to determine the bias terms for each participant. We expected to observe a stronger attraction bias when the main and secondary tasks were executed with the same hand (corresponding condition) than with different hands (non-corresponding condition). Therefore, we conducted a one-tailed paired-samples t-test between the bias of the compatible and incompatible visuomotor interference conditions. Final statistical tests of behavioral analyses were conducted in JASP (Love et al., 2019), after transferring processed data from MATLAB, unless stated otherwise.

#### Exploratory analysis

To assess the effect of different types of interference on WM representations, we compared the precision across conditions for each participant. For this analysis, errors were signed as positive if they were clockwise from the target color, and negative if they were counterclockwise. We then calculated the standard deviation (SD) of the signed errors for each condition separately for each participant and defined precision as 1/SD. As in Hypothesis II, we conducted a one-way repeated measures ANOVA with three levels (visuomotor interference, motor interference, baseline).

Furthermore, we compared response times (RT) and accuracies in the interference task. We aimed to investigate whether differences in hand correspondence conditions were related to task difficulty or complexity. Our assumption was that if no such differences existed, there would be no variation in RTs or accuracies between the corresponding and non-corresponding conditions. We compared both RTs and accuracies using a 2x2 one-way ANOVA (motor/visuomotor interference, corresponding/non-corresponding response hand).

### EEG recording, preprocessing and analyses

#### Recording

EEG was recorded with 64 Ag/AgCI passive electrodes (Easycap Gmbh, Herrsching, Germany) with a sampling rate of 1000 Hz. NeurOne Tesla AC amplifiers (Bittium Biosignals Ltd, Kuopio, Finland) were used for data collection with a 250 Hz low-pass filter. We used the FCz channel as the reference and AFz as the ground electrode position.

#### Preprocessing

We used MATLAB (R2023b, Mathworks, Natick, USA), the ERPLAB toolbox (Lopez-Calderon & Luck, 2014), and the EEGLAB toolbox (Delorme & Makeig, 2004) for the analysis of the EEG data. First, a high-pass filter with a 0.1 Hz threshold and a low-pass filter with a 40 Hz threshold were applied using an IIR Butterworth filter via the ERPLAB Toolbox’s *pop_basicfilter* (default order: 3*(sampling rate/low cut off)) function. After filtering, channels with dense artifacts were rejected using the *pop_rejchan* function (kurtosis threshold: 15), an automated process provided by EEGLAB. Frontal electrodes were excluded from this rejection procedure to maximize the capture of eye-related variance in the subsequent independent component analysis (ICA). Following channel rejection, the data were re-referenced by setting the average of all channels as the new reference. The steps between re-referencing and ICA were performed only as preparation for ICA. After calculating the IC weights, the pipeline reverted to the re-referencing stage, where the IC weights were applied to the re-referenced data, disregarding the intermediate steps.

##### ICA specific steps and IC rejection

After re-referencing, the data were downsampled to 200 Hz and high-pass filtered with a 1 Hz threshold using a Hamming-windowed sinc FIR filter (*pop_eegfiltnew,* filter order: 661, transition bandwidth: 1 Hz, cutoff frequency at −6 dB: 0.5 Hz). The data were then epoched, starting 1000ms before stimulus onset and ending 4400ms after stimulus onset. A baseline correction was applied using the 200ms period prior to stimulus onset. Automated trial rejection was then performed using EEGLAB’s *pop_autorej* function (threshold: 500 µV, maximum percentage of rejected trials per iteration: 5). ICA was applied following trial rejection. Independent components (ICs) were then identified using the ICLabel classifier for EEGLAB, classifying components into categories such as brain, muscle, eye movements, heart activity, channel noise, line noise, and “other” (Pion-Tonachini et al., 2019). ICs with a probability of being eye movement greater than 30%, and with a probability of being brain signal less than 10%, were marked for later rejection.

##### After ICA steps

After IC rejection, the remaining IC information was attached back to the re-referenced data, bypassing the ICA preparation stages. The data were then epoched from -1000ms to 4400ms, time point zero being the onset of the target stimulus. A baseline correction was applied using the 200ms pre-stimulus time window. IC components labeled as artifacts were removed. Any remaining noisy trials were rejected using the same automated trial rejection method applied during the ICA preparation stages. Finally, missing channels were interpolated using the *pop_interp* function in EEGLAB, with spherical spline interpolation, and the data were downsampled to 250 Hz.

Our preprocessing pipeline resulted in rejecting 32 ICs, on average (SD = 6.49). Excluding the blink artifacts (which are large deflections), the rejected ICs explained on average 17.82% of the variance across all participants (SD = 24.92), including components related to muscle activity, heartbeats, line noise, channel noise, and other artifacts. As a result of artifact rejection, we had an average of 933.32 remaining trials (SD = 151.51). Remaining trial counts per condition were as follows: Visuomotor corresponding (Mean = 143.42, SD = 16.06), Visuomotor non-corresponding (Mean = 144.5, SD = 16.34), Motor corresponding (Mean = 141.50, SD = 14.14), Motor non-corresponding (Mean = 142.75, SD = 17.80), and Baseline (Mean = 282.07, SD = 31.24). On average, 63.32 channels remained after preprocessing (SD = 1.38).

#### Time-frequency decomposition

We first calculated the time-frequency decomposition of the data. For each trial and channel, the oscillatory power for each frequency was calculated from 4 to 30 Hz on a logarithmic scale in 26 steps. For each frequency in the given spectrum, a complex Morlet wavelet was created by tapering a sinusoid (ei2ft) of the given frequency with a Gaussian [(e-t2/2s2; s is the width of the Gaussian; s = /(2f) corresponds to the number of cycles of wavelet)].

Our zero padded data, and the Morlet wavelets were Fast Fourier transformed (FFT). Convolution in TF decomposition was done by multiplying the signal’s Fast Fourier transform with the Fast Fourier transform of a complex Morlet wavelet in each frequency by taking the dot product. Then an inverse Fast Fourier transform was applied to get the time-domain result. We applied a baseline normalization by averaging the -400ms to -100ms time interval in all trials and divided each trial by this averaged value and converted the measurement to decibel (dB).

#### Registered analysis

To explore the first part of Hypothesis I, which predicts the onset of motor planning during the visual encoding and early maintenance phase of to-be-memorized colors, we calculated contralateral mu/beta suppression following target presentation. This refers to the suppression of ∼8-30 Hz activity on the side opposite to the hand being used for a response in the main task, measured over the centro-parietal EEG channels (C3, C4, CP3, CP4) (Rösner et al., 2022). Previous research has established the contralateral mu and beta suppression as a signature of action planning (Boettcher et al., 2021; Nasrawi et al., 2023; Rösner et al., 2022; Schneider et al., 2017; van Ede et al., 2019).

First, we calculated both contralateral and ipsilateral power across all time points and frequencies by assigning channels regarding their position to the main task response hand. Contralateral power was always derived from the channels on the side opposite to the responding hand for the main task in each trial. This analysis focused on the encoding and maintenance stage prior to interference, defined as the period from 0ms (target stimulus onset) to 1400ms (interference cue onset). Since no experimental conditions could have had an influence on EEG response in this time interval, we averaged all trials across conditions for each participant. Lateralization in oscillatory power was calculated as the contralateral-minus-ipsilateral difference. To determine whether there was significantly stronger suppression on the contralateral side, we performed a cluster-based permutation test (CBPT). This procedure began by comparing contralateral minus ipsilateral oscillatory power to zero using a one-sample t-test at each time and frequency point, identifying significant points where p < .05. We then randomly assigned the sign (positive or negative) of the difference for each participant’s data points across 1000 permutations, creating a distribution of permuted significant cluster sizes. Clusters from the original data that were larger than the 95th percentile of the permuted clusters were considered significant. For the significant cluster, we calculated Cohen’s d for each datapoint and reported the maximum and average effect size for the cluster.

Hypothesis I also predicts that greater contralateral mu/beta suppression during the encoding stage will be associated with higher WM accuracy (Boettcher et al., 2021; Nasrawi et al., 2023). To test this hypothesis, we used a median split approach for each condition separately. First, we calculated the median error value (based on absolute errors) for each participant. Then, for each condition, we calculated the average contralateral mu suppression for trials where the error was below the median and for trials where the error was above the median. To determine whether there was a significant difference in contralateral mu/beta suppression between the below-median and above-median trials, we applied the same CBPT procedure explained above. This test compared the two performance conditions over time (0-1400ms), conducted separately for each experimental condition. The aim was to identify any time-frequency clusters where a persistent and significant difference in contralateral suppression occurred between the below- and above-median trials.

#### Exploratory Analysis

While the prior analyses address motor planning processes during the encoding and maintenance phase of visual information in WM prior to interference, we also investigated to which extent the presentation of an irrelevant color affected the re-focusing on primary task information after completion of the secondary task. Prior investigations have revealed that this process can be tracked by the suppression of oscillatory power in the alpha frequency range at posterior electrodes (Woodman et al., 2022). Therefore, posterior oscillatory power was calculated as the average oscillatory power at the Oz, O1, O2, Pz, PO3, and PO4 channels. For the between-condition comparison, we applied a CBPT as explained above. The two interference conditions (motor, visuomotor) were compared against the baseline condition and against each other during the interference interval (1400-3400ms after target presentation; see Figure 3B-C). We did not average over the alpha band (8-12 Hz) to see the distribution of the effect across frequencies more clearly; however, we expected the effect to be centered around alpha.

Given that the analysis of posterior oscillatory power showed differences between the visuomotor and motor interference conditions (see Figure 3C), we aimed to examine the time course of this difference concerning the secondary task response. This approach allowed us to determine whether the onset of the difference aligned with the secondary task response, which would prove that it is rather about the refocusing on the visual content from the primary WM task than about the visual processing of the secondary task cues. To achieve this, we calculated the oscillatory power time-locked to the response to the secondary task. After the regular time-frequency decomposition, we adjusted the timeline of each trial to be centered around the response to the secondary task. Similar to the analyses of posterior oscillatory power time-locked to secondary task cue onset, we calculated the average over trials for each condition and each subject. We then performed the same comparisons as with the stimulus-onset-locked posterior alpha analysis from 700ms prior to 700ms after the response (Figure 3D; not including the baseline condition since there was no secondary task response).

In the above analyses, we conducted all examinations using pre-selected channels as regions of interest to investigate specific attentional and motor planning mechanisms. However, none of these analyses directly addressed the EEG correlates as a function of Hypothesis III (see Results), where we demonstrate the effect of hand correspondence on WM gating. Therefore, we further compared the corresponding and non-corresponding visuomotor conditions in an exploratory manner. First, we averaged the time-frequency decomposition for each condition for every participant, and we conducted another CBPT including all channels, all frequency bands and the time interval following interference presentation (1400-3400ms) across participants. To be able to explore the differences over all channels, we used the FieldTrip (Oostenveld et al., 2011) CBPT protocol. For the acquisition of the statistics, the *ft_freqstatistics* function was used (estimation method: Monte-Carlo, cluster correction, alpha: .05, t-statistics, cluster formation approach: t-sum (default), minimum number of channels for cluster formation: 2, false alarm rate: .05, number of randomizations 5000). For the calculation of neighboring channels, distances between channels were determined by the automated FieldTrip procedure. Due to the t-sum approach used in the procedure, t-values were summed for each cluster, and clusters from each permutation were distributed according to their t-sum values. The actual clusters with a t-sum higher than 95 percent of permuted clusters was considered significant.

To achieve a meaningful distribution of channel contributions to the observed effects, a procedure from Ülkü et al. (2024) was adapted. The number of significant data points from each channel was divided by the number from the channel with the highest number of significant data points, creating a value representing the relative contribution of each channel. Channels scoring more than 0.7 were considered part of the identified cluster. Significant clusters in the time-frequency domain were displayed in a contour map, along with the contributing channels.

Our exploratory analysis revealed a frontal theta difference between the corresponding and non-corresponding visuomotor interference conditions at the end of the interference cue presentation window (see Results). Further, we aimed to explore whether this TF pattern correlates with the strength of the attraction bias on a trial-by-trial basis. To calculate this, we conducted a correlation test, computing Pearson’s correlation coefficient between each time-frequency power point and the attraction bias across all trials. We then applied Fisher’s z-transformation, which involves calculating the inverse hyperbolic tangent of the coefficient distribution. This produced a matrix of time points x frequencies for each subject, each channel, and each condition (only sub-conditions of the visuomotor interference condition) with Fisher z values. We then averaged the frontal channels AF3, AF4, F1, F3 and F5 which were provided by the previous analysis as the biggest contributors of the theta effect observed (Figure 4B) and conducted a CBPT against zero for both conditions.

## RESULTS

### Behavioral results

#### Effects of interference on working memory task performance

We conducted a repeated measures ANOVA with three levels of interference type (motor interference, visuomotor interference, baseline) to determine whether adding a visual feature (the color of the interference task cue) interferes with the main task WM representation (Hypothesis II). We analyzed absolute average errors across conditions for each participant. Results indicated a main effect of interference type, *F*(2, 56) = 35.35, *p* < .001, η² = .558, BF_10_=5.48x10^8^. Post hoc comparisons revealed that visuomotor interference led to significantly higher errors compared to both the motor interference (*t*(28) = 7.551, *p* < .001, d = 0.718, 95% CI [2.933 1.488], BF_10_=6.24x10^5^) and the baseline condition (*t*(28) = 6.981, *p_holm_*< .001, d = 0.664, 95% CI [2.766 6.981], BF_10_=1.51x10^4^). No significant difference was found between the motor interference and the baseline condition (*t*(28) =0.570, *p_holm_* = .571, d = 0.054, 95% CI [0.889 -0.556], BF_10_=0.24).

We also examined the effect of corresponding vs. non-corresponding motor features between the main task and the interference task on behavioral performance to observe if shared motor features affect the handling of secondary tasks during WM storage. For this analysis, we used two factors: interference type (motor, visuomotor) and whether the response hands were corresponding or non-corresponding between tasks. To achieve this, we conducted a 2x2 repeated-measures ANOVA. Results showed a main effect of interference type (as it was shown in the previous analysis, *F*(1,28) = 59.831, *p* < .001, η² = .447, BF_10_=7.23x10^5^) and a main effect of response-hand correspondence (*F*(1,28) = 5.371, *p* =.028, η² = .04, BF_10_=2.18), with no significant interaction (*F*(1,28) = 0.018, *p* =.895, η² = 5.87x10^-6^, BF_10_=0.33). Post hoc testing indicated that when the hands used for the tasks were non-corresponding, WM accuracy was lower (*t*(28) = 2.137, *p_holm_* = .028, d= 0.207, %95 CI [0.077 1.244], BF_10_=4.4). This shows the effect of an interrupting task was slightly stronger when the two responding hands did not correspond between tasks.

Additionally, we tested whether the type of interference or the hand correspondence had an effect on the probability of using the correct hand for the main task by 2x2 repeated measures ANOVA (visuomotor/motor; corresponding/non-corresponding). Our results showed neither a main effect of interference type (*F*(1,28) = 1.417, *p* =.244, η² = .006, BF_10_=0.35), nor hand correspondence (*F*(1,28) = 2.518, *p* =.124, η² = .058, BF_10_=0.93) nor an interaction (*F*(1,28) = .204, *p* =.655, η² = .001, BF_10_=0.308).

#### Effects of interference on precision

We conducted a similar analysis on WM precision (1/SD; see methods section). The comparisons of precision yielded results consistent with those of accuracy, revealing a main effect of interference type (*F*(2, 56) = 12.915, *p* < .001, η² = 0.316, BF_10_=769.9). Specifically, precision was lower in the visuomotor interference condition than the motor interference (*t*(28) = 4.54, *p_holm_* < .001, d = 0.573, 95% CI [0.204 0.942], BF_10_=1803.5). However, precision in the motor interference condition did not differ from the baseline condition (*t*(28) = 0.299, *p_holm_* =.76, d = 0.038, 95% CI [-0.278 0.354], BF_10_=0.205).

Next, we examined the effect of hand correspondence in relation to the interference type. The analysis demonstrated a significant main effect of interference type, (*F*(1, 28) = 32.247, *p* < .001, η² = 0.252, BF_10_=2362.71). Additionally, we observed a significant effect of hand correspondence (*F*(1, 28) = 5.043, *p* =.033, η² = 0.055, BF_10_=1.725), with no significant interaction (*F*(1, 28) = 0.367, *p* =.055, η² = 0.002, BF_10_=0.28). The effect of hand correspondence suggested that precision was slightly lower when the hands did not correspond between the main task and the interference task (*t*(28) = 2.24, *p_holm_* =.033, d = 0.276, 95% CI [0.016 0.536], BF_10_=2.88).

#### Working memory gating and hand correspondence

In Hypothesis III, we predicted that shared motor features between the main and secondary tasks would trigger WM gate opening during maintenance, which should lead to an encoding of new sensory input. This will lead to an interaction between the color information already stored in WM and the task-irrelevant color of the interference task cue. Specifically, we expected this effect to manifest as an attraction bias between the interfering color and the target color in WM.

To assess whether attraction occurred, we first compared both conditions (corresponding and non-corresponding response hands) against zero using a one-tailed one sample t-test. The analysis showed that WM representations were attracted to the interfering color in both corresponding (*t*(28) = 6.936, *p<*.001, d = 1.288, 95% CI [2.427 ∞]; BF_10_= 201668) and non-corresponding conditions (*t*(28) = 6.077, *p<*.001, d = 1.128, 95% CI [1.534∞]; BF_10_= 24493).

Next, we directly compared attraction biases between corresponding and non-corresponding response hands in the visuomotor condition using a one-tailed paired-samples t-test (Figure 2C). Results indicated that when response hands corresponded between the two tasks, attraction biases were significantly higher (*t*(28) = 2.580, *p*=.008, d = 0.479, %95 [0.370∞]; BF_10_= 6.265).

#### Interference task response times and accuracy

We compared the interference task RTs using a repeated measures 2x2 ANOVA (motor, visuomotor; corresponding, non-corresponding hand) to see any possible differences in the interference task performance as a function of hand correspondence (Figure 2E). Our results showed no main effect of interference type (*F*(1, 28) = 2.526, *p* = .123, η² = .03, BF_10_=0.679) or hand correspondence (*F*(1, 28) = 0.109, *p* = .743, η² = .002, BF_10_=0.302). While there was a significant interaction (*F*(1, 28) = 5.55, *p* = .026, η² = .032, BF_10_=2.99), no post-hoc comparisons reached significance (Visuomotor interference: corresponding, non-corresponding *t*(28) = 1.509, *p_holm_*=.69, d = 0.072, %95 [-0.066 0.209]; Motor interference: corresponding, non-corresponding *t*(28) = -0.94, *p_holm_*=1, d =-0.045, %95 [-0.181 0.120]; Corresponding hands: visuomotor, motor *t*(28) = 2.654, *p_holm_*=.64, d = 0.115, %95 [-0.015 0.244]; Non-corresponding hands: visuomotor, motor *t*(28) =-0.053, *p_holm_*=1, d = -0.002, %95 [-0.124 0.120]). This makes the interpretation of the interaction rather difficult.

Furthermore, we compared the proportion of correct hand use in the interference task (Figure 2D). Our 2×2 repeated measures ANOVA revealed no significant main effects (Visuomotor, motor interference: *F*(1, 28) = 1.417, *p* = .244, η² = .006, BF_10_=0.355; Corresponding, non-corresponding hands: *F*(1, 28) = 2.518, *p* = .124, η² = .058, BF_10_=0.928) and no significant interaction (*F*(1, 28) = 0.204, *p* = .655, η² = .001, BF_10_=0.302). These results suggest that the observed difference in attraction bias between corresponding and non-corresponding conditions were not confounded by variations in task difficulty.

**Figure 2.**
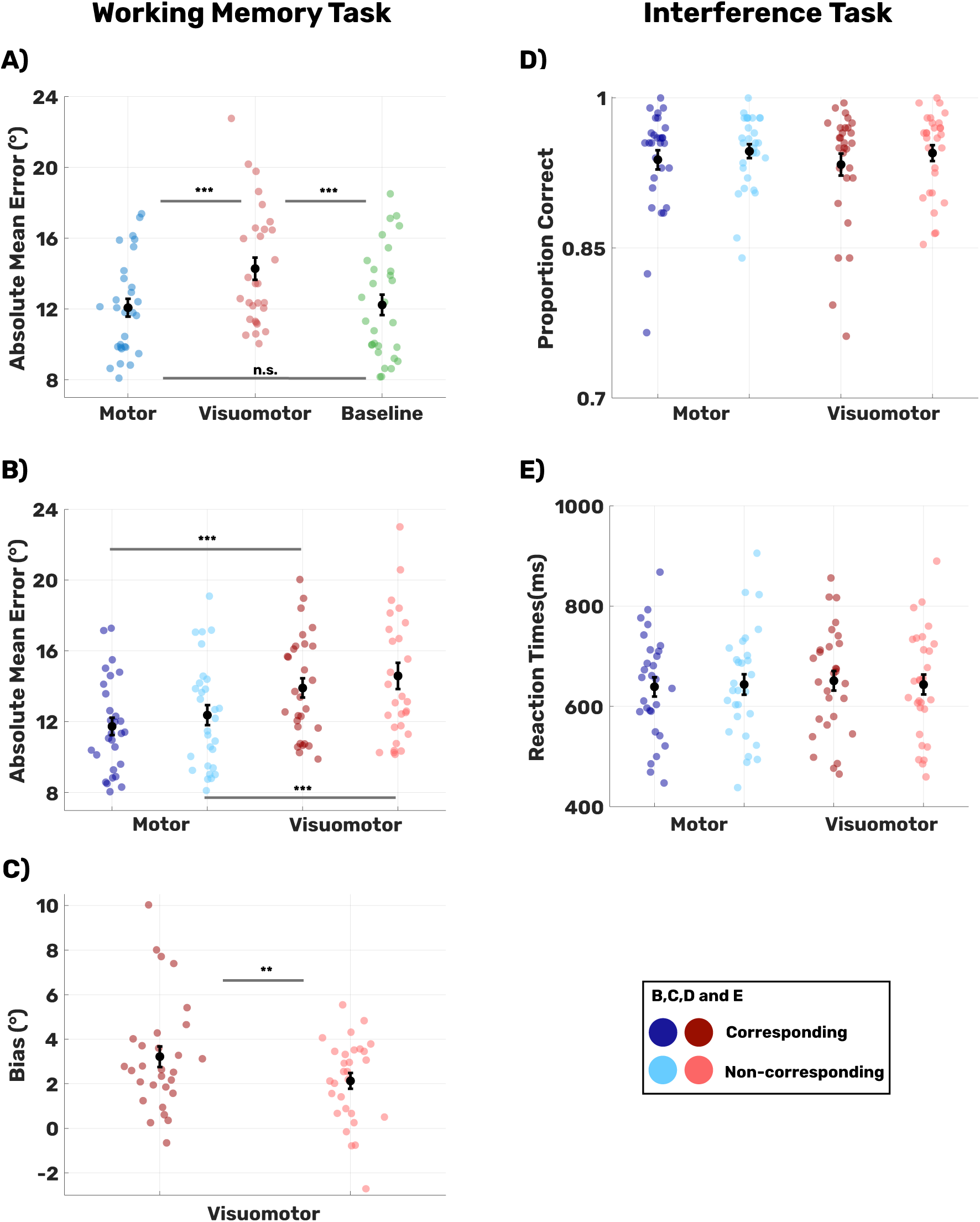
Illustration of Behavioral Performance. (*p<.05, **p<.01, ***p<.001). **A–B)** These panels compare absolute mean errors in the main task. **A)** The highest error rates occurred in the visuomotor interference condition, while motor interference did not differ from baseline. **B)** Hand manipulation had a marginal effect on main task performance. Errors were higher in the non-corresponding interference condition. **C)** The WM representation was drawn toward the interfering color (attraction bias) in both corresponding and non-corresponding conditions, with a stronger bias in the corresponding condition. **D–E)** There were no significant differences between conditions in interference task performance. Both the proportion of correct hand usage and reaction times (RTs) remained consistent across conditions.

## EEG

### Contralateral mu/beta suppression

In line with Hypothesis 1, we tested whether contralateral mu suppression occurred during the encoding of visual information into WM prior to the interference. Results showed a significant cluster in the contralateral minus ipsilateral oscillatory activity following stimulus onset, between 180-1400ms and 6.48-30 Hz (cluster size: 4426, d_max_=-1.244, d_mean_=-0.726; see Figure 3A). The suppression was distributed in the mu/beta range and was localized over centroparietal regions contralateral to the responding hand in the main task, as expected. This finding suggests that motor encoding overlaps with the encoding of sensory information.

For the second part of Hypothesis 1, we also predicted that this motor preparation would be predictive of participants’ behavioral accuracy, as shown in previous studies (Boettcher et al., 2021; Nasrawi et al., 2023). To test this, we split trials into high- and low-performance conditions based on each participant’s median accuracy score within each condition. Then, we compared averaged high- and low-performance trials using a CBPT for each condition. Our results revealed no significant clusters in any condition. Thus, although the mu/beta power lateralization before the secondary task onset reflects the encoding of the response side for the later WM report, this process did not determine the accuracy or response times (Text S3; Figure S3) of this report

### Posterior Alpha Suppression

To investigate whether the re-focusing of attention on the WM representation after an interfering task differed between the two interference conditions, we focused on differences in oscillatory activity over visual areas. We compared all conditions (motor interference, visuomotor interference, baseline) with each other, based on CBPT from the interference onset to the probe onset (1400ms – 3400ms). Results showed that posterior suppression of alpha to beta power (∼8-30 Hz) was stronger in the visuomotor interference condition than the motor interference condition (between 1668-3400ms and 4-30Hz, cluster size=6550, cluster threshold=1125.5, d_max_= -1.505, d_mean_=-0.699; see Figure 3C). Oscillatory suppression in the motor interference condition was also higher than in the baseline condition (between 1624-2572ms and 4-27.6Hz, cluster size=2637, cluster threshold=1138.5, d_max_=-1.170, d_mean_=-0.576).

We further analyzed posterior oscillatory activity in a response-locked manner, focusing on oscillatory activity time-locked to the responses to the interference task. As shown in Figure 3D, a clear difference in posterior alpha activity emerged between the motor and visuomotor conditions, largely following the interfering task response (between -248-700ms and 4-30Hz,cluster size=3758, cluster threshold=666, d_max_=-1.45, d_mean_=-0.707). This observation suggests that the posterior alpha power differences reflect refocusing on the to-be-reported WM representation, rather than differences in processing the interference task.

**Figure 3.**
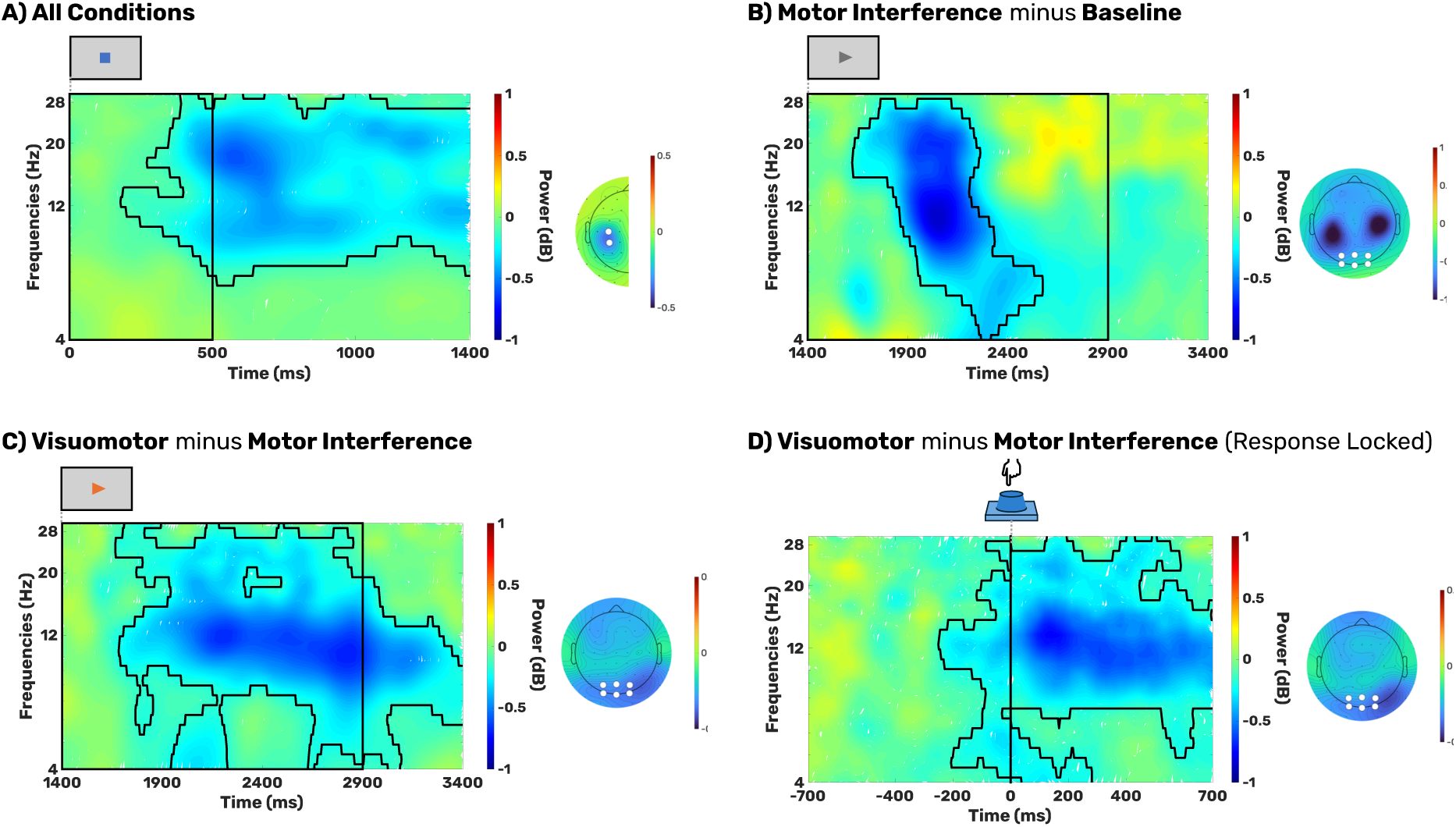
Early Motor Planning, and Reallocation of Attention After Interference. Black outlined areas indicate significant clusters identified through CBPT. White dots on each topography represent the selected channels used for the comparison. **A)** As an indication of motor planning, mu/beta lateralization (channels: C3, C4, CP3, and CP4; calculated as contra-ipsilateral channels regarding the selected hand) begins early, aligning with the target’s appearance on the screen. The topography reveals that this effect is localized to centro-parietal electrodes. The rectangle frame marks the target presentation window (0-500ms). **B)** Posterior alpha/beta suppression (averaged over Oz, O1, O2, Pz, PO3, and PO4) is higher following the motor interference when compared against the baseline condition. However, the topography shows that this difference is largest over left and right sensorimotor areas. **C)** Posterior alpha/beta suppression is most pronounced in the visuomotor interference condition. The topography shows widespread negativity across the scalp, with a stronger suppression of oscillatory power at posterior electrodes. The rectangle frame highlights the interference window (1400–2900ms). C) The posterior alpha/beta suppression difference between interference conditions emerges largely following the interference task response. The vertical line at 0ms reflects the button press in the interference task.

### Exploratory Analysis: Frontal Theta and Bias Effect

As an exploratory analysis, we investigated potential oscillatory EEG effects linked to the attraction bias differences between the corresponding vs. non-corresponding response types in the visuomotor condition (Figure 4A). To this end, we conducted an additional CBPT across times, frequencies and channels (for details, see the Methods section). This analysis revealed a difference in theta power during the later interval between the presentation of the secondary task cue and the memory probe (between 2368-2784ms and 4-8.26Hz, t_sum_=2094, t_crit_=1918, d_max_=0.75, d_mean_=0.57). Frontal theta power was higher when the hands used in the main and the interference tasks matched in the visuomotor condition. The channels contributing most to this effect were AF3, AF4, F1, F3, and F5 (Figure 4A). Notably, there was no evidence of differences in posterior oscillatory patterns between the corresponding and non-corresponding conditions. This suggests that the sensory information was processed similarly across these conditions, ruling out significant variations in early attentional orienting to the irrelevant color as the primary cause of the attraction bias effect.

To investigate the predictive relationship between the frontal theta difference and the attraction bias effect toward the secondary task cue color, we calculated single-trial correlations between time-frequency (TF) data and bias scores for each time point, frequency, and channel. A CBPT was then conducted on these correlation scores, restricted to the frontal channels identified in the prior exploratory analysis. This effect was examined separately for the corresponding and non-corresponding response conditions.

Our analysis revealed that only in the visuomotor corresponding response condition did frontal theta power predict bias scores at the single-trial level (Figure 4B). Specifically, a negative relationship was observed: higher frontal theta power was associated with a lower attraction bias (between 2636-3056ms and 4-7.62Hz, cluster size=689, cluster threshold=489, dmax=-1.09, dmean=-0.61). These results suggest that the increase in frontal theta power reflects a reactive control mechanism for mitigating secondary task interference.

**Figure 4.**
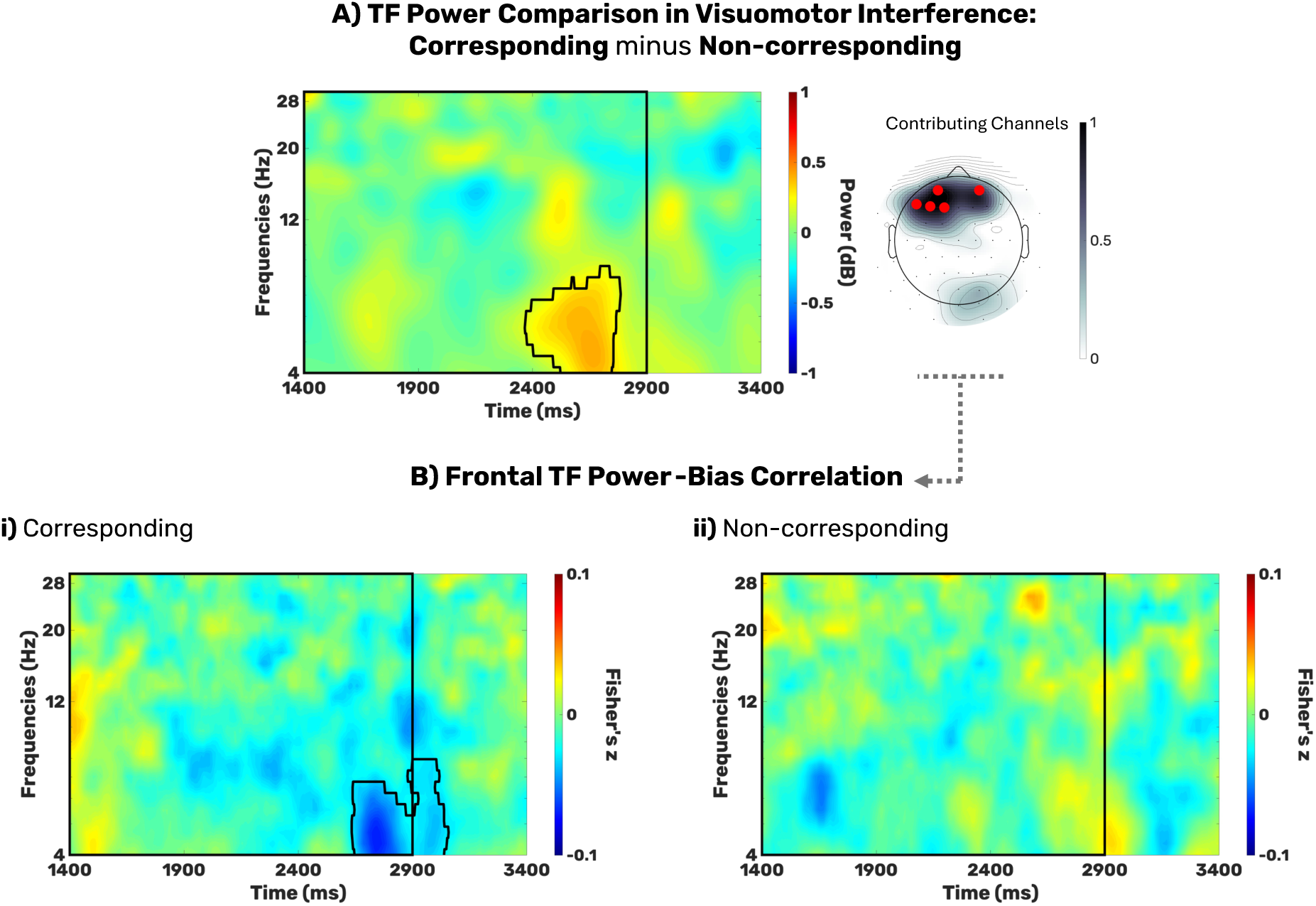
Response-Related Bias Effect and Frontal Theta Power. Outlined areas in the contour plots represent significant clusters identified through CBPT. Black rectangle frames represent the interference window (1400ms-2900ms). **A)** An exploratory analysis reveals increased frontal theta activity in the late phase of the interference interval in the corresponding visuomotor interference condition compared to the non-corresponding condition. The marked channels of the topography plot highlight the key contributing channels (AF3, AF4, F1, F3, and F5). **B)** In the corresponding visuomotor condition only, frontal theta activity (calculated over the contributing channels from Figure 4A) predicts the level of attraction bias on a trial-by-trial basis at the end of the interference interval. Higher theta power is associated with lower attraction bias. The time-frequency distribution of this effect closely resembles the frontal theta differences observed between conditions.

## DISCUSSION

In this study, we tested the role of motor processes in WM gating. Our results showed that when the planned motor responses for a WM task matched the response required for an interference task, the irrelevant visual information presented during the interference task influenced later WM report to a stronger extent. This finding suggests that motor control processes can modulate the extent to which WM is prone to irrelevant sensory information.

First, there was a main effect of interference type on absolute errors in the WM task. As shown in Figure 2A, the increase of errors was linked to the presentation of irrelevant sensory information (visuomotor condition). There was no general increase in errors by responding to a secondary task during WM storage (motor condition). This supports the notion that irrelevant visual information during maintenance can disturb visual WM representations (Lorenc et al., 2021; Rademaker et al., 2015; Saito et al., 2023), at least when this information is presented within a sensory object requiring a response. We further observed an effect of hand correspondence on absolute errors in the WM task (see Figure 2B). When the interference task was done by the non-corresponding hand, the magnitude of absolute errors increased. This might indicate that using different hands for the two tasks creates conflict and increased cognitive demand, impairing the task performance.

In line with our main hypothesis, our results further revealed higher levels of attraction bias to the interfering color during visuomotor interference when the responding hand corresponded between the two tasks (see Figure 1C). It is important to note that these two behavioral parameters (i.e., absolute angular error and signed errors / the bias) exhibited opposite effects of hand correspondence. This supports their different conceptualization: the increase of absolute errors with non-corresponding responses reflects a drop of overall task (i.e., memory) performance, whereas the increase in the attraction bias reflects a stronger representational shift toward the interfering color. This modulation of the attraction bias aligns with the view that WM input gating can be influenced by actions (Chatham & Badre, 2015). According to this view, such gating adaptations may allow WM to respond dynamically to task-relevant environmental changes during action execution. This process is thought to be mediated by the basal ganglia (BG), which play a dual role in motor control (Cui et al., 2013) and sensory gating mechanisms (Slagter et al., 2012). While our study does not provide region/network-specific evidence for this mechanism, our findings conceptually align with the framework of BG-mediated WM gating. This means that when the execution of a certain motor action corresponds to motor plans already stored in WM, it may lead to the uptake of sensory information into WM resulting in attraction bias. However, it is also possible that the mere requirement to respond to the cues in the visuomotor interference task leads to a transfer of the interference color into WM, and the representational shift towards the interference color occurs at a later stage. In this scenario, the response correspondence effect would reflect a later-stage effect during memory storage.

Since we calculated the bias parameter as the average of signed errors based on their position relative to the interference color, it was impossible to determine whether the effect was primarily driven by the bias in WM or by swap errors (i.e., misreporting the interference color instead of the target). To address this, we compared two WM models (Text S1): Target Confusability Competition Model (TCC; Schurgin et al., 2020) and TCC Swap Model (Williams et al., 2022); and found that the swap-free model provided a better fit to error distributions across participants (Figure S1A). Furthermore, when we estimated a bias parameter using the TCC model (Saito et al., 2025), the results replicated an higher attraction bias in the corresponding hand condition (Figure S1B). Together, these findings support our interpretation of the effect as an attraction bias rather than being driven by swap errors.

In order to break down the underlying processes in more detail, we further explored how the neural oscillatory patterns are linked to the effect of motor correspondence on WM bias. There was higher frontal theta power for the corresponding than for the non-corresponding visuomotor interference condition at the end of the interference task interval. This frontal theta power predicted the level of attraction bias only in the corresponding visuomotor interference condition, with higher theta power associated with a lower attraction bias (see Figure 4B). Given the role of frontal theta power in conflict resolution (Cohen & Donner, 2013; Kaiser et al., 2022; Nigbur et al., 2011), we interpret this activity as reflecting a mechanism that actively mitigates interference arising from action-induced opening of the WM gate. The fact that the relationship between theta power and bias in WM is exclusively observed in the corresponding condition (Figure 4B) further demonstrates the increased reliance on top-down control only when the response hands match between the tasks. This interpretation aligns with findings by Rac-Lubashevsky and Kessler (2015; 2018), who demonstrated that when there is an automatic tendency to update WM, but the task requires maintaining existing information, higher theta activity is observed. It is further plausible that this control of interference is related to the focusing attention back to WM (de Vries et al., 2018; Gresch et al., 2025; Zickerick et al., 2021). This assumption is supported by the fact that other oscillatory parameters such as the modulations on posterior alpha power (Figure 3D) and the lateralization of the mu/beta power (Text S4; Figure S4.) also occur in the same time window. While these oscillatory effects reflect different processes, they might have in common that they are temporally linked to the resumption of the WM task.

Given that the motor plan interacted with the attraction effect by the visuomotor task, it must have been initiated before the interference task. In line with this, mu/beta suppression contralateral to the responding hand emerged during the presentation of the to-be-memorized object (Figure 3A). While some studies have suggested that such early correlates of motor planning can predict precision and response times of the WM task (Boettcher et al., 2021; Nasrawi et al., 2023), our findings did not support this notion by neither showing an effect on accuracy nor response times. However, our study did not allow participants to form precise action plans due to the randomization of the color wheel on each trial. While participants knew which hand to use, they could not anticipate the exact degree of rotation required. In contrast, Nasrawi et al. (2023) and Boettcher et al. (2021) allowed for a better prediction of the motor requirements during target report, because the starting orientation of the memory probe was fixed, enabling participants to plan the specific amount of rotation required. Additionally in terms of response times, participants may have prioritized accuracy over speed given the long response window.

Furthermore, we observed differences in posterior alpha/beta activity (∼8-30 Hz) between interference conditions during the secondary task period (Figure 3C). Most notably, suppression of posterior alpha/beta power was stronger in the visuomotor interference condition than in the motor interference condition. This difference appeared largely after the interference task response (Figure 3D), suggesting that it emerged after the processing of the interfering task cue was complete (i.e., when no further sensory processing of the interfering task cue was required). The stronger suppression of alpha/beta power after the visuomotor interference could therefore indicate greater demands for refocusing attention on the visual information stored in WM. Moreover, our results show that the timing of the re-emergence of mu/beta lateralization corresponds with the response times in the secondary task (Figure S4). These results are in line with Gresch et al. (2025) showing a timely re-orienting of attention to WM following the completion of an interrupting task.

## Limitations

One limitation of our study is the absence of a visual interference condition without any action requirement. Including such a condition could have clarified whether the observed drop in memory performance in the visuomotor condition was driven purely by irrelevant sensory input or depended on a required motor response. However, the main focus of the current investigation was the interaction between motor processes and sensory interference in WM.

Moreover, the additional condition might have allowed us to assess whether the hand correspondence effect regarding the increased attraction in the corresponding condition or reduced attraction in the non-corresponding condition (or both). Nevertheless, our finding that theta power correlated with attraction bias exclusively in the corresponding condition aligns with previous reports of increased cognitive control demands during automatic, task-detrimental WM updating (Rac-Lubashevsky & Kessler, 2015, 2018).

## Conclusion

In this study, we demonstrated that motor processes influence WM susceptibility to interference, with corresponding motor demands increasing attraction towards task-irrelevant information. To our knowledge, this study provides the first experimental evidence for motor processes affecting WM gating, observed at both behavioral and EEG levels. Additionally, we proposed that the uptake of irrelevant information into WM is counteracted by a reactive control mechanism reflected in frontal theta oscillatory activity, aimed at preserving the storage of task-relevant information in WM.

## Data and Code Availability

All data, materials, scripts and experimental setup are shared publicly on Open Science Framework (https://osf.io/7fve8/). Only anonymized data are shared.

## Preregistration

The experimental design, data collection plan, hypotheses, and analysis plan for this study have been preregistered on the Open Science Framework prior to data collection (https://doi.org/10.17605/OSF.IO/CBZY2).

## Author Contributions

All authors contributed to the conceptualization of this study. Ş.Ö. and D.S. designed the experiment. Ş.Ö. collected the data. Ş.Ö. and D.S. conducted the data analysis. All authors discussed the interpretation of the results and wrote the manuscript.

## Competing Interests

The authors declare no competing interests.

## Notes

https://osf.io/7fve8/

### Competing Interest Statement

The authors have declared no competing interest.

### Summary of Updates

The current version includes supplemental material that provides additional analyses for a more comprehensive interpretation.

