## Supplemental Material for "Motor control processes moderate visual working memory gating"

### Text S1: The models and model fitting

To determine whether our bias analysis is influenced by swap errors, which means mistakenly reporting the interference color, we conducted additional modeling analyses. First, we compared the Target Confusability Competition (TCC) Model (Schurgin et al., 2020) with the TCC Swap Model (Williams et al., 2022). The models are explained in detail below; however, the main difference is that the TCC Swap model estimates the frequency of swap errors in the data. We selected this comparison because other swap models in the literature assume equal memory strength for the target and interference items. In our study, the interference color is not part of the memoranda, making it unlikely that the target and interference share the same memory strength. Given this important aspect, we used the TCC Swap which modifies the model to estimate different memory strengths for the interference and memory items. We compared these two models using Akaike Information Criterion (AIC) and Bayesian Information Criterion (BIC). For model fits and model comparisons, we used the Maximum Likelihood function of MemToolbox (Suchow et al., 2013). As explained below, the model comparison indicated that our data's error distribution is better explained by the TCC model without the swap error component.

#### Target Confusability Competition Model

As a signal detection-based theory of working memory (WM), the Target Confusability Competition Model assumes that responses in a memory task are based on a familiarity signal that is, participants select the item that feels most familiar at the time of response.

The model posits that each color on the color wheel has an associated memory signal determined by its psychophysical similarity to the target. On each trial, the model assumes that the participant chooses the option with the strongest signal.

More formally, the similarity function is denoted as $\mathrm{f}(x)$, and $d'$ represents the memory strength of the target. The memory signal for each option on the color wheel is drawn from a normal distribution with a mean of $d' \times f(x)$ and unit variance. The final response corresponds to the index with the maximum signal value across all colors. As the familiarity function, we used a similarity function in color space previously derived from a familiarity judgment task (for details, see Schurgin et al., 2020). This can be expressed as:

$$r \sim argmax(X_{-179},...,X_{180})$$

**Target Confusability Competition + Swap Model**

TCC Swap model is a modification of the model structure explained above. TCC swap model first proposed by Williams et al., (2022) to estimate frequency of swap errors.

The modified model estimated four parameters: $\theta, \beta, d'T,$ and $d'P.$ In the model, $\beta$ represents the proportion of swap errors (ranging from 0 to 1); and $d'T$ and $d'P$ correspond to the memory strengths for the target and the distractor. Similar to previous memory signal for each option on the color wheel is drawn from a normal distribution with a mean of $d' \times f(x)$ and unit variance, and they allow let $(Y - 179, \ldots, Y180)$ represent the probe distribution with means $dy = d'P * g(y)$ and unit variance for distribution of distractor selection when there is a swap. Then the response can be modelled as:

$$\omega \sim Bernoulli(\beta)$$
$$r \sim \omega \times argmax(Y\_\{-179\},...,Y\_\{180\}) + (1-\omega) \times argmax(X\_\{-179\},...,X\_\{180\})$$

**Model Analyses Results**

We compared the goodness of fit measures of two proposed models on the error distribution of the visuomotor interference condition. Since we have visual interference only in the visuomotor condition, only this condition allowed us to observe possible swap errors or bias in the distribution. We compared individual fits by comparing the AIC and BIC scores using paired sample t-tests. However, since AIC and BIC scores deviate normality, we also report the Wilcoxon test results as a non-parametric alternative. Our results showed that the TCC model was a better fit, supported by both AIC ($t(28) = -6.092, p < .001, d = -1.131, BF_{10} = 12704; z=$ 4.703, p<.001), and BIC ($t(28) = -18.629, p <.001, d =-3.459, BF_{10}= 1915 \times 10^{12}$; z = 4.703, p<.001), as indicated in Figure S1A. Lower AIC/BIC scores indicate better fits.

We conducted the same analysis on signed errors (as explained in the main text, Methods section, Registered Analysis: Hypothesis III) by assigning all distractor positions as positive and still obtained the same results for AIC ($t(28) = -4.862, p < .001, d = -0.903, BF_{10} = 592.763$; $z= -4.011$, p<.001) and BIC ($t(28) = -15.920, p < .001, d = -2.956, BF_{10} = 4.03 \times 10^{12}; z= -4.703$, p<.001). This indicates that swap errors with the interference color do not explain the error distributions of visuomotor interference condition.

Then, to further examine our bias comparison results from the main text, we implemented a bias parameter using the WithBias() function of MemToolbox (Suchow et al., 2013), allowing the TCC model's estimated distribution to shift (Saito et al., 2025). This way, we estimated the shift in the distribution of errors (parameter $\mu$) besides the memory strength. We fitted the model to the signed error distributions of each individual in the visuomotor corresponding-hand and visuomotor non-corresponding-hand conditions. A positive estimate of $\mu$ indicated that the distribution is shifted toward the interference color, as errors were assigned negative or positive relative to the interference color's position.

Our model estimates showed the same pattern as the model-free approach (Figure S1B): the estimated bias was positive and significantly above zero for both conditions (corresponding: $t(28) = 8.140$, $p < .001$, $d = 1.512$, $BF_{10} = 1.737x10^6$; non-corresponding: $t(28) = 4=5.406$, $p < .001$, $d = 1.004$, $BF_{10} = 2303$ ), and it was higher in the corresponding-hand condition ($t(28) = 3.808$, $p < .001$, $d = .707$, $BF_{10} = 45.705$).

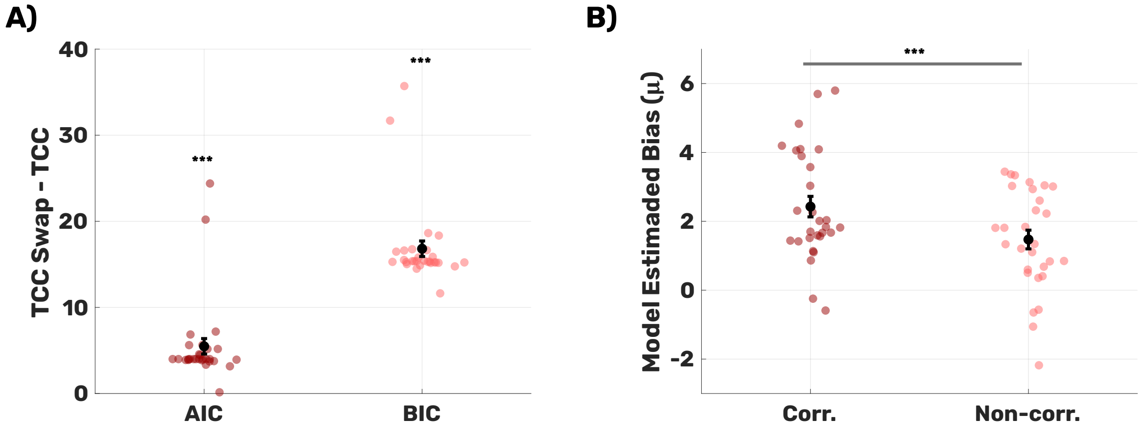


**Figure S1. Model Comparisons and Model-based Bias Estimations.** (*$p$<.05, **$p$<.01, ***$p$<.001) **A)** Both AIC and BIC comparisons across participants suggested that the TCC model without swaps fit our error distribution better, indicating that our error distribution was not explained by swap errors. **B)** Model-based bias estimations showed the same pattern of bias: both hand correspondence conditions exhibited attraction towards the interference color, with a stronger effect observed in the corresponding hand condition.

### Text S2: Event-related potentials (ERPs)

As explained in the Discussion section, we wanted to see whether there is a surprise-related effect in the visuomotor interference condition due to the changing colors of the visuomotor cue. This would be possible if, in certain trials, a randomly selected color attracts attention because it represents a rare stimulus within the overall experimental procedure. Specifically, we explored whether this difference between the motor (only grey cues) and visuomotor conditions (variable cue color) induces an oddball effect in the event-related potential (ERP; Polich & Margala, 1997; Reed et al., 2022).

To investigate this, we compared the ERPs at electrode Pz across both conditions. ERPs were calculated as the activity at Pz following standard preprocessing steps (including baseline removal). For each subject, trials were averaged per condition, resulting in time-locked amplitude data per condition. We then compared these data using a cluster-based permutation test (see Methods section of the main text for CBPT details). Our analysis did not show any difference in the ERPs between the visuomotor and motor interference conditions (Figure S2). Since both conditions involved the same motor requirements but differed in their visual features, we interpret these results to suggest that the interference effects we observed cannot be driven by a surprise-related effect resulting from the changing cue colors.

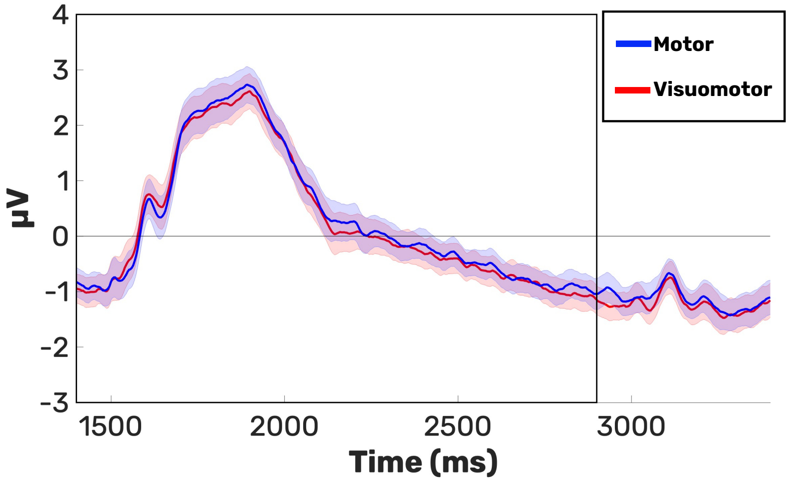


**Figure S2. ERP Comparison Between Visuomotor and Motor Interference.** The figure demonstrates the Pz ERPs following the interference onset (1400ms; the black square indicates the interference period). The shaded spaces show the standard errors for each time point. Our CBPT analysis showed no difference at any time point between the visuomotor and motor interference ERPs. This suggests that the changing colors in the visuomotor interference condition did not trigger a surprise effect.

### Text S3: Response times and mu/beta lateralization

As discussed in the Discussion section, recent studies have shown that stronger mu/beta lateralization regarding the response hand during the encoding stage is associated with faster response times during report (Boettcher et al., 2021; Nasrawi et al., 2023). To test whether our data reflect a similar pattern, we calculated the median response times of the main task for each participant and categorized trials as either above or below this median RT per subject and experimental condition. RTs were calculated as the onset of the first movement of the response knob. We then averaged the contralateral minus ipsilateral time-frequency data per subject (for details on TF calculation and electrode selection see Methods section, Analyses, Time Frequency), and compared these differences between the above- and below-median RT trials using a CBPT. Our results showed no significant cluster in this comparison (Figure S3 A;B). We further interpret these results in the discussion section.

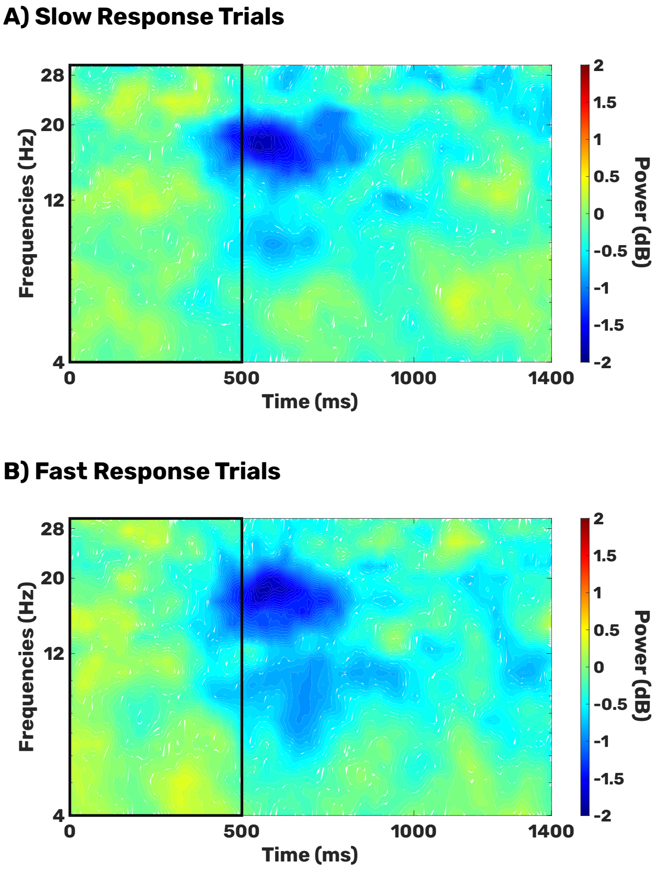


**Figure S3. Mu/Beta Lateralization Comparison Between Slow and Fast Response trials.** Figures show contra-minus-ipsilateral time-frequency (TF) decomposition over C3, C4, CP3, and CP4 channels relative to the response hand. The 0–500 ms box indicates the stimulus presentation window. **A)** Contralateral TF suppression for slow response trials, calculated from trials with response times above the subject-specific and condition-specific medians for the memory task. **B)** Contralateral TF suppression for fast response trials, calculated from trials with response times below the subject-specific and condition-specific medians for the memory task. Our CBPT comparison revealed no significant difference between the fast and slow trials.

#### Text S4: Discontinuation of mu/beta lateralization by the interference

Our data supported the idea by Gresch et al. (2025) and Zickerick et al. (2021)that attention is redirected to the primary task immediately after having responded to an interrupting task. We further explored whether response hand selection for the main task is prioritized immediately following the interference task. To test this idea, we compared the contralateral mu/beta suppression for the visuomotor and motor interference conditions combined across the entire trial period. As Figure S4A shows, during the interference period, mu/beta suppression is interrupted and then recovers (cluster size: 12699, $d_{max} = 1.737$, $d_{mean} = .803$).The onset time of this recovery corresponds to the response time of the interference task (Mean = 644.49ms, SD = 105.068ms), as indicated by the red line in Figure S4A. This indicates that once the interference task is done, the motor codes of the main task are prioritized immediately. Furthermore, we conducted the same comparison time-locked to the secondary task response (Figure S4B). In line with our previous findings, this analysis showed that the re-emergence of motor planning for the main task was time-locked to the secondary task response following the interruption (cluster size: 3870, $d_{max} = 1.609$, $d_{mean} = .742$).

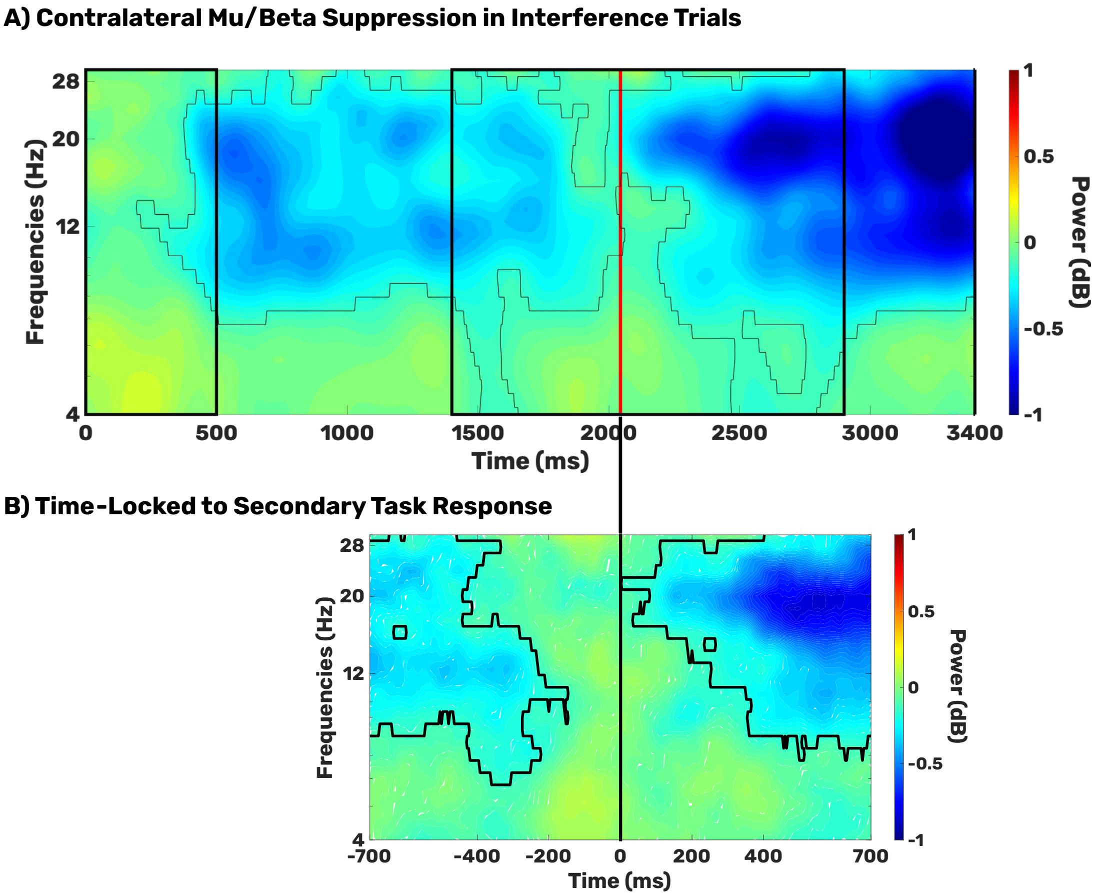

**Figure S4. Contralateral Mu/Beta Suppression; Visuomotor and Motor Interference**. The black box on the left represents the stimulus presentation window (0-500ms), while the larger black box on the right indicates the interference time window (1400-2900ms). Marked areas with black lines show the significant clusters. A) The red line on the figure represents the average RT of the interfering task. Recovery of the mu/beta lateralization following the interference corresponds to the average response time of the interference task. B) Black line (time point 0) indicates the secondary task response. The response-locked analysis shows the re-emergence of the contralateral mu/beta suppression is time-locked to the secondary task response.
